# Complete Genome Sequence of the *Wolbachia w*AlbB Endosymbiont of *Aedes albopictus*

**DOI:** 10.1101/415521

**Authors:** Amit Sinha, Zhiru Li, Luo Sun, Clotilde K. S. Carlow

## Abstract

*Wolbachia*, an alpha-proteobacterium closely related to Rickettsia is a maternally transmitted, intracellular symbiont of arthropods and nematodes. *Aedes albopictus* mosquitoes are naturally infected with *Wolbachia* strains *w*AlbA and *w*AlbB. Cell line Aa23 established from *Ae. albopictus* embryos retains only *w*AlbB and is a key model to study host-endosymbiont interactions. We have assembled the complete circular genome of *w*AlbB from the Aa23 cell line using long-read PacBio sequencing at 500X median coverage. The assembled circular chromosome is 1.48 megabases in size, an increase of more than 300 kb over the published draft *w*AlbB genome. The annotation of the genome identified 1,205 protein coding genes, 34 tRNA, 3 rRNA, 1 tmRNA and 3 other ncRNA loci. The long reads enabled sequencing over complex repeat regions which are difficult to resolve with short-read sequencing. Thirteen percent of the genome is comprised of IS elements distributed throughout the genome, some of which cause pseudogenization. Prophage WO genes encoding some essential components of phage particle assembly are missing, while the remainder are scattered around the genome. Orthology analysis identified a core proteome of 536 orthogroups across all completed *Wolbachia* genomes. The majority of proteins could be annotated using Pfam and eggNOG analyses, including ankyrins and components of the T4SS. KEGG analysis revealed the absence of 5 genes in *w*AlbB which are present in other *Wolbachia*. The availability of a complete circular chromosome from *w*AlbB will enable further biochemical, molecular and genetic analyses on this strain and related *Wolbachia*.

**Data deposition:** Raw data from PacBio sequencing have been deposited in the NCBI SRA database under BioProject accession number PRJNA454708, as runs SRR7784284, SRR7784285, SRR7784286, SRR7784287. The paired-end reads from Illumina library used for indel correction are available from NCBI SRA database as accession SRR7623731. The assembled genome and annotations have been submitted to the NCBI GenBank database with accession number CP031221.

## Introduction

*Wolbachia* are gram-negative α-proteobacteria of the order *Rickettsiales*. Maternally transmitted infections are widespread, occurring in an estimated 40-65% of insect species (Hilgenboecker et al. 2008; Werren et al. 2008) including 28% of mosquito species (Kittayapong et al. 2000). *Wolbachia* infections are not limited to arthropods as nematodes, including several major human pathogens, also harbor the endosymbiont (Fenn et al. 2006; Lefoulon et al. 2016). Currently, *Wolbachia* strains have been classified as *Wolbachia pipientis* (Hertig 1936; Lo et al. 2007), which has been divided into 16 major phylogenetic clades termed supergroups, denoted A-Q, mainly on the basis of 16S rDNA phylogenetic analyses. Most supergroups are restricted to arthropods (A, B, E, G, H, I, K, M, N, O, P and Q) (Lefoulon et al. 2016), whereas supergroups C and D are the major nematode-infecting lineages. Supergroup F is unique as it contains both nematode and arthropod-infecting strains (Lefoulon et al. 2016). The nature of the association between *Wolbachia* strains and their hosts varies greatly. In nematodes, the prevalence of infection is 100% and the relationship is obligate (Taylor et al. 2005). These attributes have been exploited to enable the use of antibiotics as a novel approach to treat filarial infections (Langworthy et al. 2000; Bazzocchi et al. 2008; Johnston et al. 2014). In contrast, infection is less prevalent in insect hosts and can cause broad effects on insect physiology leading to several phenotypic changes attributed to the ability of *Wolbachia* to act as manipulators of the host (Werren et al. 2008; Cordaux et al. 2011). Among these manipulations, cytoplasmic incompatibility (CI) is the most common phenotype in mosquitoes (Sinkins 2004) and provides a reproductive advantage to *Wolbachia*-infected females over uninfected females, resulting in spread and persistence of *Wolbachia* in populations (Xi et al. 2005). When experimentally transferred to uninfected mosquitoes, *Wolbachia* can also suppress infection or transmission of viruses (Walker et al. 2011; Aliota, Walker, et al. 2016; Aliota, Peinado, et al. 2016; Carrington et al. 2018), *Plasmodium* parasites (Kambris et al. 2010) and filarial nematodes (Kambris et al. 2009; Andrews et al. 2012) making *Wolbachia* a particularly attractive agent for control of vector-borne pathogens.

The Asian tiger mosquito, *Aedes albopictus*, is an aggressive biting mosquito and currently one of the most invasive species in the world. Originally native to Southeast Asia, the species has spread in the past 30-40 years and colonized five continents (Kotsakiozi et al. 2017). It is a significant public health concern as it is a competent vector of several arboviruses that cause severe diseases in humans such as dengue, chikungunya and zika (Gratz 2004; Chouin-Carneiro et al. 2016; Grard et al. 2014). Two distinct *Wolbachia* strains (*w*AlbA and *w*AlbB), are present in variable density in *Ae. albopictus* tissues (Kittayapong et al. 2000; Zouache et al. 2009). *w*AlbB, belonging to the supergroup B, is a particularly interesting strain to study since it has been reported to induce opposing phenotypes in different hosts following either malaria or viral infection. Transient somatic infection of *Anopheles gambiae* with *w*AlbB inhibits *Plasmodium falciparum* but enhances *Plasmodium berghei* parasites (Hughes et al. 2011, 2012). It enhances West Nile virus infection in the mosquito *Culex tarsalis* (Dodson et al. 2014), whereas it blocks transmission of dengue (Mousson et al. 2012) and chikungunya (Raquin et al. 2015). The interplay between *w*AlbB and its host is also particularly important as it impacts the stability of *w*AlbB following its introduction into new hosts such as *Aedes aegypti* mosquitoes to control dengue and zika transmission to humans (Pan et al. 2018).

Cell lines containing *Wolbachia* represent a simplified model in which to explore the symbiotic relationship and have been used extensively in molecular, biochemical and genetic studies (O’Neill et al. 1997; Voronin et al. 2012; Saucereau et al. 2017). The Aa23 cell line derived from *Wolbachia*-infected *Ae. albopictus* mosquito embryos was the first cell line developed to enable studies on *Wolbachia*-host cell interactions (Sinkins et al. 1995; O’Neill et al. 1997). While *Ae. albopictus* mosquitoes are naturally infected with *w*AlbA and *w*AlbB, only *w*AlbB was retained in the Aa23 cell line (Sinkins et al. 1995; O’Neill et al. 1997). The cell line comprises at least two cell types and *Wolbachia* infection varies, with respect to both the level of infection among individual cells and the overall level of infection in a population (O’Neill et al. 1997). However, high cell density during passaging helps to maintain a relatively stable infection rate, because the duration of exponential growth is affected by cell density (Gerenday & Fallon 1996). *w*AlbB from Aa23 cells has been used as a source of infection for other insect cell lines (Dobson et al. 2002; Fenollar, Scola, et al. 2003; Xi et al. 2005; Rasgon et al. 2006). Since no nematode-derived cell culture system for *Wolbachia* exists, the Aa23 insect cell line:*w*AlbB model system has been used as a proxy to screen for new anti-*Wolbachia*/filarial compounds (Fenollar, Maurin, et al. 2003).

Due to the importance of the *w*AlbB-infected Aa23 cell line in studies on symbiosis, pathogen control and drug screening, a draft genome sequence of this strain was published (Mavingui et al. 2012). For this *Wolbachia* assembly, Multiple Displacement Amplification of DNA from infected cells was used to construct a mate-paired library containing 6-kb inserts, and sequenced with 454 Titanium pyrosequencing at 76 bp read-length. The resulting genome draft is incomplete with 165 contigs encompassing 49 scaffolds (Mavingui et al. 2012), hampering a comprehensive analysis of the genome.

The short-read technologies, such as 454 and Illumina, cannot easily reconstruct complete microbial chromosomes, and often produce draft assemblies containing gaps. Pacific Biosciences (PacBio) SMRT technology produces long reads, some as long as 100 kb, with average raw read lengths >15 kb, making single and continuous assembly possible (Eid et al. 2008). In addition, the PacBio library preparation process does not include an amplification step, therefore DNA is sequenced as a single molecule in its native form, enabling the detection of covalent base modifications (Flusberg et al. 2010).

In the present study, we have assembled the complete circular genome of *w*AlbB present in the Aa23 cell line, from long-read PacBio sequencing data at 500X median coverage. The long reads enabled sequencing over complex repeat regions which have been difficult to resolve with short-read sequencing. The assembled circular genome is 1,484,007 bp in size, an increase of 321 kb over the published *w*AlbB draft genome, making it one of the largest sequenced *Wolbachia* genomes to date. This sequence will serve as important resource for detailed studies of *w*AlbB and related *Wolbachia*.

## Materials and Methods

### Cell culture

The Aa23 cell line infected with *w*AlbB was a kind gift from Dr. Stephen Dobson. Cells were grown in culture flasks at 28 °C in equal volumes of Mitsuhashi–Maramorosch medium (Sigma M9257) and Schneider’s insect medium (Sigma S0146), supplemented with 10% heat-inactivated fetal bovine serum (O’Neill et al. 1997). The cells retained the morphological heterogeneity originally described (O’Neill et al. 1997) and were routinely sub-cultured at 7 to 10 day intervals by diluting 1:3 in fresh media to maintain high cell density and contiguous monolayers.

### Immunostaining of *Wolbachia* with anti-VirB8 antibody

Cells cultured on glass coverslips within 24-well microtiter plates were fixed in 4% formaldehyde in phosphate-buffered saline (PBS) for 15 min and subsequently permeabilized using chilled 100% methanol (−20°C) for 1 min. Fixed cells were then incubated in polyclonal rabbit anti–VirB8 antibody (Li & Carlow 2012) diluted 1:2,000 in PBS containing 5% goat serum, followed by Alexa Fluor 488 (green) conjugated goat anti-rabbit secondary antibodies (Molecular Probes; Invitrogen Life Technologies) according to manufacturer’s instructions. Cell nuclei were stained with Hoechst 33342 at 1:10,000 dilution in PBS. Prolong Gold anti-fade reagent (Invitrogen Life Technologies) was used to avoid fading. Images were acquired using an Axiovert 200M microscope (Carl Zeiss, Oberkochen, Germany) and processed using ZEN software (Carl Zeiss).

### DNA extraction

To harvest host cell-free *w*AlbB, spent culture media from Aa23 cells (passage #65) was collected and centrifuged at 500g to remove cell debris, followed by 5,000g to collect the *Wolbachia*-enriched pellet. Genomic DNA was extracted using a Qiagen MagAttract HMW kit following manufacturer’s instructions. Briefly, 220uL of buffer ATL and 20uL of proteinase K were added, and the sample was incubated at 56°C for 3 hours with mixing at 900 rpm (Eppendorf thermomixer C). DNA was eluted with 150uL of AE buffer and quantified using a NanoDrop and Qubit instruments (Thermo Fisher Scientific). The quality of DNA was assessed using an Agilent 4200 TapeStation System. The DNA obtained was good quality (DIN > 8.2), and high molecular weight, larger than 60 kb in size (Supplementary Figure S1).

### PacBio and Illumina library construction and sequencing

For library construction, intact genomic DNA was fragmented using a Megaruptor 2 device (Diagenode). Sheared DNA was purified with AMPure PB beads and 2µg were used to construct a SMRTbell library according to PacBio library construction guidelines with some modifications. Briefly, sheared DNA was repaired using the NEBNext FFPE DNA Repair Mix, followed by end-repair to generate blunt ends. Following purification using AMPure PB beads, PacBio universal hairpin adaptors were ligated to the DNA to produce SMRTbell libraries. After adaptor removal and library clean up, concentration and size of the SMRTbell library were determined using the Qubit HS DNA kit and Agilent TapeStation analysis. To enrich for longer insert sizes, size selection was performed using the BluePippin system (Sage Science), resulting in a library that contained an insert size of approximately 45 kb (Supplementary Figure S1). The PacBio sequencing primer was then annealed to the SMRTbell library followed by binding of the polymerase to the primer-library complex. The size-selected library was loaded onto 2 SMRT cells in the PacBio RSII system using a MagBead binding kit and sequenced with a 300 minutes collection time. Two additional SMRT cells were loaded with library that did not undergo size selection.

For Illumina library construction, genomic DNA was fragmented to 300 bp average size using a Covaris S2 (Covaris Inc.) with the following settings: 10% duty cycle, intensity 4,200 cycles per burst and treatment time of 80 seconds. Libraries were constructed using the NEBNext Ultra II DNA Library Prep Kit for Illumina (New England Biolabs, Inc.). The library quality was assessed using a high sensitivity DNA chip on a Bioanalyzer (Agilent Technologies, Inc.). The library was sequenced on an Illumina MiSeq platform (paired-end, 150 nt reads).

### Genome Assembly and DNA modification analysis

PacBio sequencing reads from all 4 flow cells were processed and assembled using the HGAP assembler version 3 (Chin et al. 2013) as implemented in the PacBio SMRT^®^ Analysis Server v2.3.0 (https://www.pacb.com/products-and-services/analytical-software/smrt-analysis). The contig corresponding to *w*AlbB was selected for further polishing and circularization. Overlapping regions at the termini of this contig were identified by BLAST analysis, and were merged to circularize the chromosome. This circular draft assembly sequence was further polished using multiple rounds of the ReSequencing.1 protocol from the PacBio SMRT^®^ Analysis Server v2.3.0. The origin of replication, *oriC*, was identified by generating a consensus of all 10 *Wolbachia oriC* sequences obtained from the DoriC database (Gao et al. 2013). The assembled chromosome was verified to be free of any structural errors via the RS.Bridgemapper pipeline available as a part of the PacBio SMRT portal.

The validity and correctness of chromosome circularization was confirmed by PCR and sequencing across the ends of the polished chromosome.

Primers F1 (5’TCCCCTGCCCTACCTGAGTA3’) and R1 (5’GTCATCATCCTGCGCGAGAG3’) were used to amplify a 1,599 bp fragment that spans the junction of circularization; primers F2 (5’ TGTTGCTTTCATTGAGGCTGGT3’) and R2 (5’ TATTGGACCCACACCGCGAA3’) were used to amplify a 1081bp fragment to verify the *oriC* sequence, using the Q5 HiFi PCR master mix (NEB M0543) following manufacturer’s instructions. Search for potential DNA modifications in the *Wolbachia w*AlbB genome was carried out using the polished genome as a reference genome in the RS_Modification_and_Motif_Analysis.1 pipeline from the PacBio SMRT^®^ Analysis Server v2.3.0.

To check and correct any potential indel errors typically observed in PacBio-only assemblies (Watson 2018), Illumina sequencing was performed. After adapter-trimming and filtering of low quality reads using BBMap package, version 37.17 (https://sourceforge.net/projects/bbmap), the reads were mapped to the PacBio chromosome using bwa version 0.7.15-r1140 (Li & Durbin 2009) in paired-end mode. Pilon software version 1.22 (Walker et al. 2014) was run on the bam file output from bwa, using alignments with mapping quality ≥ 20 (Pilon flag minmq=20).

### Genome annotation and analysis

Protein-coding genes, rRNA, tRNA, ncRNA and pseudogenes were identified using the NCBI prokaryotic annotation pipeline (Angiuoli et al. 2008). Further functional annotation of protein-coding genes was carried out using the eggNOG-Mapper (Huerta-Cepas et al. 2017) web server (http://eggnogdb.embl.de/#/app/emapper) against the eggNOG database (Huerta-Cepas et al. 2016). The completeness of the genome was assessed using the BUSCO pipeline version 3.0.2 (Simão et al. 2015). Insertion sequence (IS) elements were identified by searching against the ISfinder database (Siguier et al. 2006) via the ISsaga web server (Varani et al. 2011) available at http://issaga.biotoul.fr/issaga_index.php. Pfam domains were annotated using the pfam_scan.pl script version 1.6 from http://ftp.ebi.ac.uk/pub/databases/Pfam/Tools to search against Pfam database version 31.0 (Finn et al. 2016). Annotation of integrated prophage regions was carried out using the PHASTER web server (Arndt et al. 2016), available at http://phaster.ca, and by comparisons to other *Wolbachia* prophage sequences. These include WOVitA1 (GenBank: HQ906662.1), WOCauB2 (GenBank: AB478515.1) and WOCauB3 (GenBank: AB478516.1), WOVitB (GenBank: HQ906665.1, HQ906666.1) and the prophage regions from *w*Mel (GenBank: NC_002978.6). Circos plots (Krzywinski et al. 2009) for visualizing the distribution of various features across the genome were plotted using the R package circlize, version 0.4.3 (Gu et al. 2014). Search for orthologs across multiple genomes was performed using the OrthoFinder (Emms & Kelly 2015) software version 1.1.4. The number of orthogroups common across various *Wolbachia* were visualized as UpSet plots (Lex et al. 2014) using the R package UpSetR, version 1.3.3 (Conway et al. 2017).

KEGG automatic annotation server, KAAS, (Moriya et al. 2007), available online at https://www.genome.jp/kegg/kaas, was used to find functional annotations of genes in the *w*AlbB genome. *w*AlbB protein sequences were used as query sequences and blast (bi-directional best hit) searched against a manually curated set of ortholog groups in KEGG to generate KEGG pathways and functional classifications. The KO assignments of *w*AlbB proteins from KEGG pathway analysis were then compared to the KEGG pathways of *Wolbachia w*Ri from *Drosophila simulans* and *w*Pip from *Culex quinquefasciatus* available in the KEGG database, to identify any missing proteins in *w*AlbB.

## Results

### High levels of *Wolbachia* infection enable production of host cell-free *Wolbachia*

To preserve high levels of *Wolbachia* infection in Aa23 cells, maintenance of a high-density monolayer of cells was found to be necessary, which was achieved by passaging at high cell densities. Approximately 80% of cells were infected with a high *Wolbachia* load as verified by staining with an α-virB8 antibody (Figure 1A) or a Hoechst 33342 DNA dye (Figure 1B). The large numbers of host cell-free *Wolbachia* observed in spent culture media (Figure 1C), obviated the need for further separation of *Wolbachia* from host cells.

### Assembly and Annotation

Processing of the combined PacBio data obtained from 4 SMRT cells produced 944,546 filtered subreads and a total of 3 billion bases, with the longest subread at 61 kb and median length 3.4 kb. HGAP3 assembly of this data generated 581 contigs. The longest contig was 1,511,710 bp, at ∼500X median coverage (Supplementary Figure S1) and 99.9994% consensus concordance. This single contig contained all the contig sequences from the published *w*AlbB assembly (RefSeq assembly accession GCF_000242415.2), and was selected for further polishing. No other contig matched the *w*AlbB genome. One contig at 2,400X median coverage corresponded to *Ae. albopictus* mitochondrial DNA, and the remaining contigs mostly matched *Ae. albopictus* genomic regions.

**Fig. 1.**
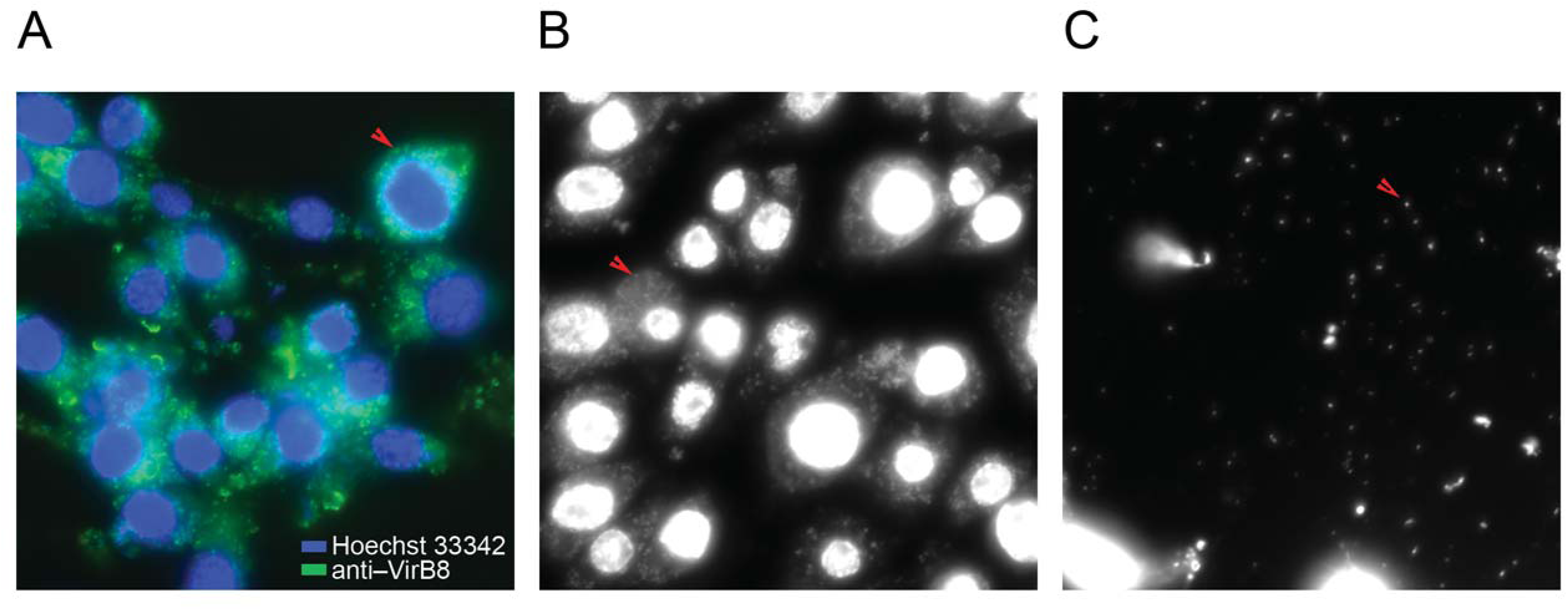
Detection of *Wolbachia* in Aa23 cells and in culture supernatants. Immunostaining of *Wolbachia* using an anti-VirB8 antibody (green) and Hoechst staining (blue) of host nuclei (A). Hoechst staining of *Wolbachia* in cells (B) and in spent media (C). Arrows indicate *Wolbachia.*

A BLAST analysis of the *w*AlbB contig to itself identified a highly similar 27 kb region repeated at the beginning and end of the contig, indicating that this contig represents the complete circular chromosome. To validate that these regions (marked A1 and A2 in Figure 2A) represent overlapping ends of the circular chromosome, we collapsed them into a single consensus region (marked A in Figure 2A-B). Using the PacBio ReSequencing.1 pipeline, all sequencing reads were re-mapped to this corrected chromosome sequence, which generated a polished single contig representing a circular chromosome with non-repeating ends. Primers F1 and R1 (Figure 2A-B) designed to span this candidate junction of circularization produced a PCR product of expected size of 1.5 kb (Figure 2C) and sequence, confirming the correctness of the circularization. For the next round of polishing, the first base of the chromosome sequence was reset to the start of *oriC* and this permuted sequence was again used as a reference for re-mapping all the reads using the PacBio ReSequencing.1 pipeline. The sequence of the *oriC* region was also verified by sequencing the PCR product generated using primers F2 and R2 (Figure 2A-C). The final polished circular genome produced as the output had a median coverage of ∼500X and was used in all subsequent analysis. The identified *w*AlbB *oriC* has all the hallmark features typical of *Wolbachia oriC* regions (Ioannidis et al. 2007). It is flanked by *hemE* and *tlyC* genes, with the intergenic regions having binding sites for DnaA, IHF and CtrA (Figure 2D).

**Fig. 2.**
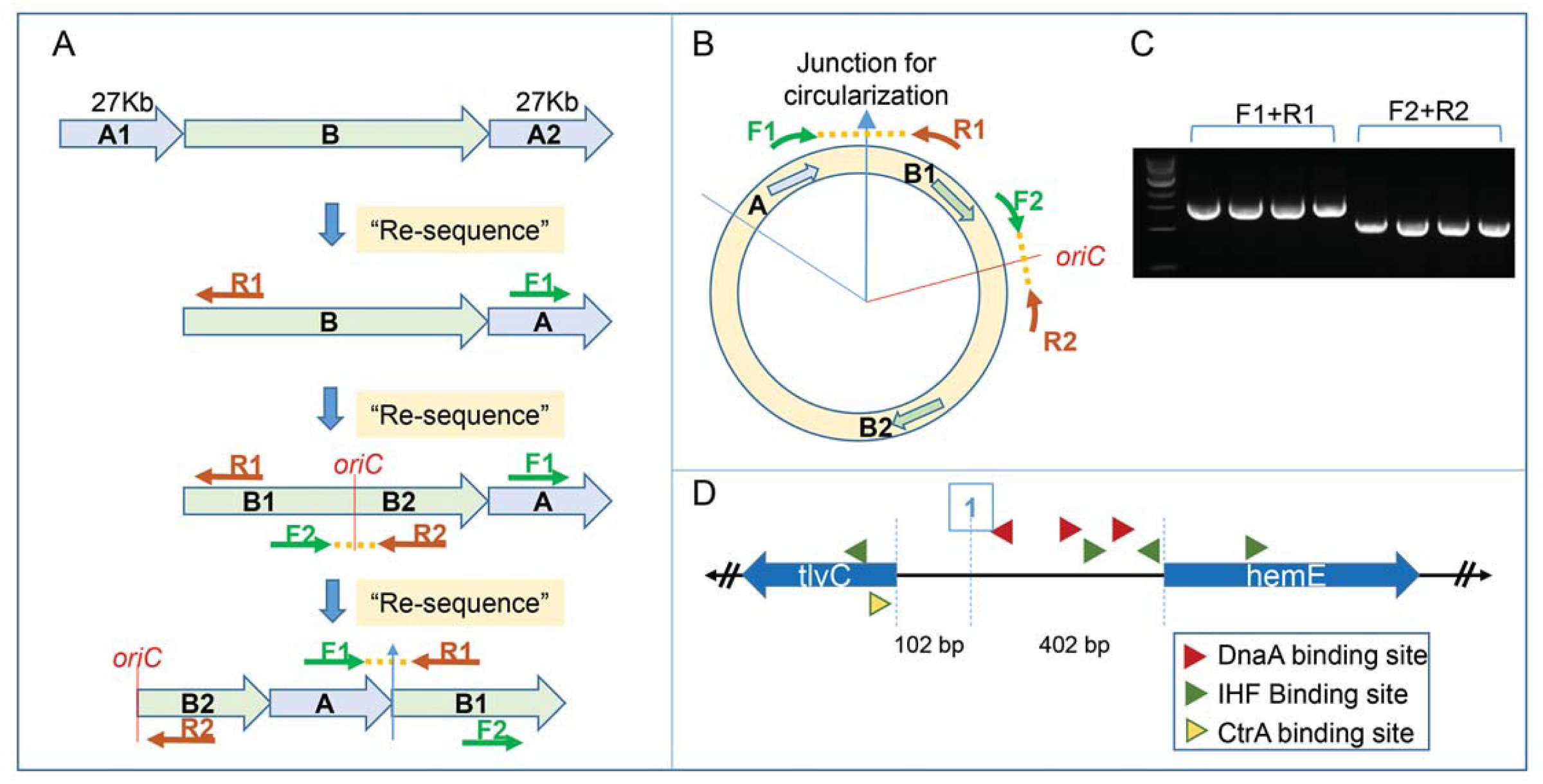
Circularization and polishing of the *w*AlbB genome assembly and determination of *oriC*. *De novo* assembly of PacBio data produced a single contig representing the *w*AlbB genome, with terminal regions marked A1 and A2 in (A), showing high sequence identity over a 27 kb region, suggesting overlapping ends of a circular chromosome. They were therefore collapsed into a single consensus region “A” to represent the circular the chromosome, with the junction of circularization between regions A and B indicated by a blue arrow (B). This draft circular genome was polished using the PacBio ReSequencing.1 pipeline, and the junction of circularization was validated by Sanger sequencing of a 1.5 kb amplicon (C) produced by primers F1 and R1. The origin of the circular chromosome was reset to the beginning of the *oriC* locus. The permuted chromosome sequence was again polished using ReSequencing.1 pipeline in SMRT analysis software. The *oriC* sequence and the new junction of circularization at *oriC* was verified by primers F2 and R2 (A and B) that successfully produced an amplicon of correct size 1.5kb (C) and correct sequence as confirmed by Sanger sequencing. The *oriC* locus in *w*AlbB has all the hallmarks observed in other *Wolbachia oriC* sequences, such as flanking genes *tlyC* and *hemE*, three DnaA binding sites, four IHF binding sites and one CtrA binding site (D). All PCR were performed on four independent DNA samples.

Indel errors are sometimes observed in assemblies produced from only long-read technologies such as PacBio or Nanopore (Watson 2018). Therefore, the polished, circular *w*AlbB genome was checked for any potential errors using Illumina reads. About 4.8 million paired-end reads mapped to the assembly (mapping quality ≥ 10) providing a ∼630X median coverage of the genome and were used as an input to the Pilon error-detection and correction tool (Walker et al. 2014). This analysis did not identify any indels, but 65 potential single nucleotide polymorphisms (SNPs). However, these SNPs have very low quality scores and low coverage (∼2X coverage as opposed to a median coverage of ∼630X at all other positions) and therefore did not pass the filter for making changes. In summary, no errors were detected in the PacBio assembly based on the Pilon analysis using high-quality and high coverage Illumina read alignments. This indicates that the median PacBio coverage of ∼500X and multiple rounds of polishing using the PacBio ReSequencing.1 pipeline was sufficient to produce an accurate assembly.

The final circular chromosome is 1,484,007 bp in size (Table 1), an increase of 321 kb over the published *w*AlbB draft genome. Using the nucmer tool for genome alignments (Kurtz et al. 2004), all the contigs from the published genome (Mavingui et al. 2012) could be mapped to the complete assembly (Figure 3A). The regions common between the complete genome and the previously published draft assembly share > 99% sequence identity. The average GC content of the genome is 34.4 %, which is within the typical range for *Wolbachia* genomes (Table 1). Analysis with RS_Modification_and_Motif_Analysis.1 pipeline did not identify DNA modifications such as m4C or m6A suggesting that these modifications are absent from the *w*AlbB genome (Figure S2).

**Table 1.** Key characteristics of all complete *Wolbachia* genomes.

| Table 1. Key characteristics of all complete *Wolbachia* genomes |  |  |  |  |  |  |  |  |  |  |  |  |  |
| --- | --- | --- | --- | --- | --- | --- | --- | --- | --- | --- | --- | --- | --- |
| Name | Host organism | Supergroup | RefSeq assembly accession | Size (Mb) | GC% | Proteins | rRNA | tRNA | Other RNA | Total Genes | Pseudogenes (%) | Ankyrin proteins | BUSCO score |
| w Au | *Drosophila simulans* | A | GCF_000953315.1 | 1.26 | 35.2 | 1,099 | 3 | 34 | 4 | 1,265 | 125 (9.9) | 21 | 83.8 |
| w Ha | *Drosophila simulans* | A | GCF_000376605.1 | 1.29 | 35.3 | 1,126 | 3 | 35 | 4 | 1,263 | 95 (7.5) | 26 | 83.3 |
| w Mel | *Drosophila melanogaster* | A | GCF_000008025.1 | 1.27 | 35.2 | 1,100 | 3 | 34 | 4 | 1,270 | 129 (10.2) | 19 | 82.9 |
| w Ri | *Drosophila simulans* | A | GCF_000022285.1 | 1.45 | 35.2 | 1,254 | 3 | 34 | 4 | 1,403 | 108 (7.7) | 25 | 83.3 |
| w AlbB | *Aedes albopictus* (Aa23 cell line) | B | GenBank Accession = CP031221 | 1.48 | 34.4 | 1,205 | 3 | 34 | 4 | 1,434 | 188 (13.1) | 34 | 81.9 |
| w No | *Drosophila simulans* | B | GCF_000376585.1 | 1.30 | 34.0 | 1,065 | 3 | 34 | 4 | 1,231 | 125 (10.2) | 41 | 82.4 |
| w Pip | *Culex quinquefasciatus* | B | GCF_000073005.1 | 1.48 | 34.2 | 1,257 | 3 | 34 | 4 | 1,402 | 104 (7.4) | 43 | 81.9 |
| w Tpre | *Trichogramma pretiosum* | B | GCF_001439985.1 | 1.13 | 33.9 | 827 | 3 | 35 | 4 | 1,106 | 237 (21.4) | 10 | 82.4 |
| w Oo | *Onchocerca ochengi* | C | GCF_000306885.1 | 0.96 | 32.1 | 651 | 3 | 34 | 4 | 759 | 67 (8.8) | 1 | 74.7 |
| w Ov | *Onchocerca volvulus* | C | GCF_000530755.1 | 0.96 | 32.1 | 649 | 3 | 34 | 4 | 763 | 73 (9.6) | 1 | 75.6 |
| w Bm | *Brugia malayi* | D | GCF_000008385.1 | 1.08 | 34.2 | 839 | 3 | 34 | 4 | 1,047 | 167 (16.0) | 12 | 80.6 |
| w Fol | *Folsomia candida* | E | GCF_001931755.1 | 1.80 | 34.8 | 1,513 | 3 | 35 | 4 | 1,658 | 103 (6.2) | 83 | 81.0 |
| w Cle | *Cimex lectularius* | F | GCF_000829315.1 | 1.25 | 36.3 | 981 | 3 | 34 | 4 | 1,246 | 224 (18.0) | 33 | 80.6 |

Note. - For a standardized analysis across all genomes, the number of ankyrin proteins in each genome was determined in this study, using an identical pipeline based on Pfam domain annotations. Similarly, the BUSCO scores (reported as % complete BUSCOs) for each of the genomes was also calculated using identical parameters (-lineage = proteobacteria\_odb9, 221 BUSCO groups). The other genome characteristics were obtained from the RefSeq database using the accession numbers indicated.

**Fig. 3.**
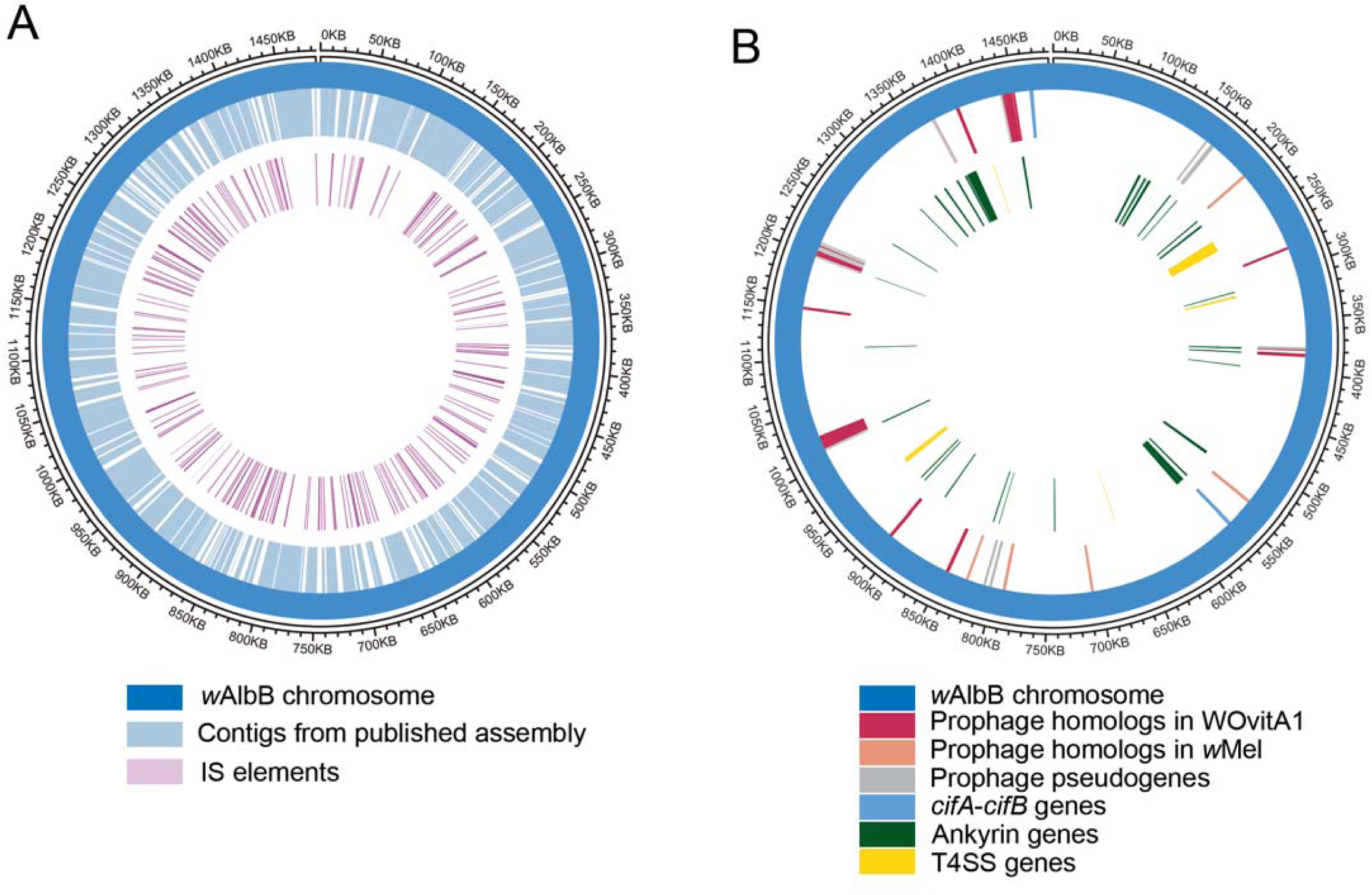
Circos plot representation of features on the circular *w*AlbB genome. The *w*AlbB genome is represented as the outer blue circle, with the coordinates marked on the outermost circle. (A) The completed *w*AlbB genome contains all the contigs from the published draft genome (depicted in light blue on the 1^st^ inner circle) revealing the gaps (white regions, 1^st^ inner circle). The IS elements (purple, innermost circle) are distributed all over the genome, and tend to be located near the gaps, close to the termini of the contigs from the published draft. (B) Positions of prophage genes and pseudogenes with orthologs in WOVitA1 and *w*Mel genomes are indicated on the first inner circle. The positions of the cifA-cifB gene pairs are also indicated on this track. Locations of genes encoding ankyrin proteins (green) and T4SS components (yellow) are indicated on the second inner circle.

Annotation of the genome using the NCBI prokaryotic annotation pipeline (Angiuoli et al. 2008) identified 1,205 protein-coding genes, an increase of 250 genes over the published version. The genome also encodes 34 tRNA genes that include cognates for all amino acids, 3 rRNA (16S, 23S, and 5S), plus another 3 non-coding RNAs (6S RNA, RNAse P RNA, and a signal recognition particle sRNA small type) and one tmRNA gene. A total of 188 pseudogenes were identified resulting from one or more of the following causes: frameshift (n = 39), incomplete (n = 150), internal stop (n = 23) or multiple problems (n = 21). The percentage of pseudogenes is comparable across all the completed *Wolbachia* genomes from various supergroups (Table 1).

The completeness of genome annotation was evaluated using the Benchmarking Universal Single-Copy Orthologs (BUSCO) pipeline (Simão et al. 2015), which measures the proportion of expected gene content from highly conserved, single-copy orthologs (BUSCO groups). The analysis was carried out against 221 BUSCO groups derived from 1,520 proteobacterial species. The 1,205 protein coding genes in the *w*AlbB genome contain 179 complete and single copy BUSCO groups, 2 complete and duplicated BUSCO groups, 6 fragmented BUSCOs and 34 missing BUSCOs, resulting in a 81% BUSCO completeness score. For comparison, the BUSCO scores were also calculated for the other completed *Wolbachia* genomes (Figure S3) and similar completeness scores were obtained for all genomes analyzed (Table 1).

### Insertion Sequence (IS) elements comprise 13% of the *w*AlbB genome

Insertion sequences are one of the simplest transposable elements, usually encoding only a transposase flanked by short direct-or inverted-repeats, and can play a major role in genome evolution even in a short time scale (Siguier et al. 2015). IS elements have been classified into ∼20 families based on sequence similarities (Siguier et al. 2006). *Wolbachia* genomes often harbor numerous IS elements and their identification is essential for a comprehensive study of *Wolbachia* genome evolution (Cerveau et al. 2011). To annotate IS elements in the *w*AlbB genome, the ISsaga web service (Varani et al. 2011) was used to query the ISfinder database (Siguier et al. 2006) and 218 IS elements were found distributed throughout the genome (Figure 3A, Table S3), including 45 partial IS elements containing pseudogenized transposase (Table S3). The IS982 and IS481 family IS elements were the most abundant, with 96 and 75 copies respectively. The median size for IS elements is 873 bp, and they range in length from 66 bp to 1,683 bp, adding up to total of 191,182 bp or about 13% of the entire *w*AlbB genome. Mapping the contigs from the published draft genome (Mavingui et al. 2012) to the completed genome revealed that break points and/or gaps overlap with IS elements (Figure 3A). This indicates that the repetitive nature of IS elements hinder genome assembly using short-read data, and this problem can be overcome by using long-read technologies such as PacBio.

Movement of IS elements can cause insertions/deletions in a genome, sometimes leading to pseudogene formation. For example, the published contig NZ_CAGB01000139.1 (17, 533 bp) was found to map to two regions of ∼540 bp and ∼17,000 bp on the complete genome, with a 1,203 bp insertion caused by an IS481 element, resulting in the formation of 2 pseudogenes (DEJ70_01295, DEJ70_01305) derived from the *rsmD* gene. PCR amplification and Sanger sequencing across this region confirmed this insertion and pseudogenization of the *rsmD* gene.

### Orthology analysis and identification of a core proteome across completed *Wolbachia* genomes

Orthology relationships between the *w*AlbB proteins and the RefSeq protein sequences from all other complete *Wolbachia* genomes (Table 1) were analyzed using OrthoFinder program, which identifies families of homologous proteins and assigns them to orthogroups (Emms & Kelly 2015). A total of 1,604 orthogroups were derived from a combined set of 13,002 proteins (Table S4). Of these, 1,171 orthogroups comprised of 12,569 proteins are shared by 2 or more genomes (“shared orthogroups”), while the remaining 433 orthogroups are unique to each *Wolbachia* analyzed (Table S4). For *w*AlbB, 1,192 of the 1,205 protein-coding genes (99%) were assigned to 918 shared orthogroups, while 13 *w*AlbB proteins could not be assigned to any such group (Table S4). Similarly, for all the other genomes analyzed, more than 93.5% of proteins could be assigned to a shared orthogroup, and the proportion of potentially genome-specific orthogroups was found to be low (up to 6.5%). The only outlier was *w*Fol, with 14% of its 1,403 proteins not be assigned to any shared orthogroup (Table S4).

The core proteome, defined as the set of proteins present in all genomes analyzed, consists of 536 orthogroups (Figure 4), and 519 of these orthogroups contain single-copy 1:1 orthologs (Table S4). Outside the core proteome, the number of shared orthologous groups decreased substantially (Figure 4).

**Fig. 4.**
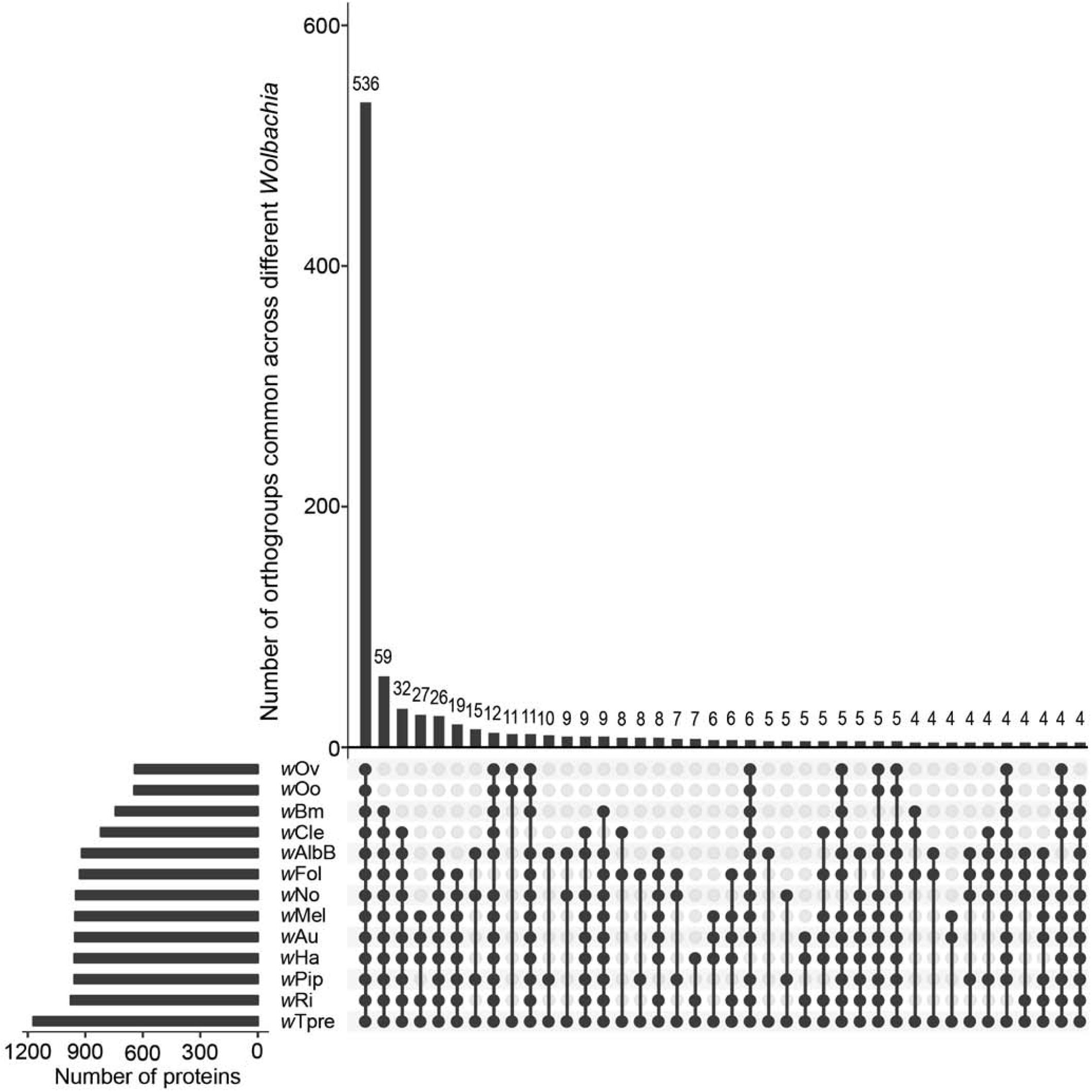
Orthology analysis of proteins from all complete *Wolbachia* genomes identifies core *Wolbachia* proteome. The set of all 13,002 proteins from *w*AlbB and 12 other completed *Wolbachia* genomes were grouped into 1,171 orthogroups using the OrthoFinder software. Of these, 536 orthogroups were present across all the genomes analyzed, representing the core *Wolbachia* proteome, represented by the first bar in the Upset plot. Other orthogroups showed various patterns of distribution. Filled dots (black) denote presence and empty dots (grey) indicate absence of orthogroups in each *Wolbachia*. Genomes of *Wolbachia* from *Drosophila melanogaster* (*w*Mel), *Drosophila simulans* (*w*Ri, *w*Ha, *w*No, *w*Au), *Culex quinquefasciatus* (*w*Pip), *Brugia malayi* (*w*Bm), *Onchocerca ochengi* (*w*Oo), *Onchocerca volvulus* (*w*Ov), *Folsomia candida* (*w*Fol), *Trichogramma pretiosum* (*w*Tpre) and *Cimex lectularius* (*w*Cle) were included in the analysis.

### Pfam and eggNOG annotations

Analysis of Pfam protein domains (Finn et al. 2016) encoded in the *w*AlbB genome identified 995 genes encoding proteins containing at least one Pfam domain, representing 83% of the total genes present (Table S1). By far the most abundant domains arise from mobile genetic elements. For example, the DDE Transposase domain (“DDE_Tnp_1_3”, Pfam accession PF13612) was the most abundant domain, present in 82 proteins, followed by the “Retroviral Integrase” domain (“rve”, Pfam accession PF00665) present in 67 proteins. We also found 48 proteins containing the reverse transcriptase domain (“RVT_1”, Pfam accession PF00078), and 53 proteins with the Group II intron reverse transcriptase domain (“GIIM”, Pfam accession PF08388). Further annotation of gene function based on orthology using the eggNOG software (Huerta-Cepas et al. 2017) could assign a putative function to 1,044 (87%) of the 1,205 *w*AlbB protein-coding genes, with transposase, integrase, and reverse transcriptase functions again being the most abundant classes.

### *w*AlbB genome contains degenerated WO prophage

Prophages play an important role in *Wolbachia* biology e.g. the prophage WO from the *Wolbachia* strain *w*Mel contributes to the cytoplasmic incompatibility in its *Drosophila* host (Masui et al. 2000, 2001; LePage et al. 2017; Beckmann et al. 2017). Availability of a complete genome made the search for any potential prophages in *w*AlbB feasible. The PHASTER webserver (Arndt et al. 2016) identified two prophage derived regions in the *w*AlbB genome. The larger prophage region is 15.4 kb in size and showed highest nucleotide similarity to the WOVitA1 prophage. Further BLAST and OrthoFinder analysis identified *w*AlbB orthologs for 40 of the 63 WOVitA1 genes, and 7 pseudogenes corresponding to 5 other WOVitA1 genes, while 18 WOVitA1 genes were found to be completely absent in the *w*AlbB genome (Table S5). The prophage genes absent in *w*AlbB include the components essential for a phage particle assembly, such as tail subunit I, baseplate subunits J, W and V, phage portal protein and phage minor capsid protein, suggesting an inactive prophage. Further, the prophage derived genes (Figure 3B, Table S5) are located in 3 separate clusters in the *w*AlbB genome containing 14, 16 and 8 genes respectively, while the remaining prophage genes are distributed over the genome. In addition, 5 genes and 6 pseudogenes that are more similar in sequence to prophage genes in *w*Mel rather than WOVitA1 were identified (Table S5). None of these additional 5 genes can compensate for functions missing in WOVitA1 derived prophage regions. Overall, the combined size of all prophage derived regions in *w*AlbB genome is 49.8 kb, comprising 3.3 % of the entire genome. Together these observations suggest that the prophages in *w*AlbB have undergone degeneration and are not active.

### CI genes in *w*AlbB

The *w*AlbB *Wolbachia* is known to cause cytoplasmic incompatibility (CI) in its *Ae. albopictus* host (Dobson et al. 2001). Genetic studies of CI in *Drosophila* hosts have identified a pair of genes, *cifA* and *cifB* (LePage et al. 2017; Beckmann et al. 2017), which are sometimes located within the WO prophage regions (Lindsey et al. 2018). A phylogenetic analysis of *cifA* and *cifB* across all *Wolbachia* has found them to co-occur as a pair of neighboring genes, and grouped them into four and three monophyletic “Types” respectively (LePage et al. 2017; Lindsey et al. 2018). A search for *cifA* and *cifB* homologs in *w*AlbB identified two sets of *cifA* and *cifB* gene-pairs. The first pair is composed of a Type IV *cifA* (DEJ70_02760) and a Type III *cifB* (DEJ70_02755), while the second pair is composed of a Type III *cifA* (DEJ70_07090) and a Type III *cifB* (DEJ70_07095). Interestingly, neither of these gene pairs are located in the prophage derived regions in *w*AlbB (Figure 3B), suggesting that they do not always need to be encoded in a prophage and can possibly be integrated into *Wolbachia* genomes.

### Type IV secretion system in *w*AlbB

Many symbiotic and pathogenic intracellular bacteria use a type IV secretion system (T4SS) for successful infection, proliferation and persistence within hosts. It is a diverse and versatile transporter system which spans the entire cell envelope functioning in conjugation, competence and effector molecule (DNA and/or protein) translocation (Grohmann et al. 2018). Genome analysis of *w*AlbB revealed the presence of a T4SS with 15 components organized in two operons and 4 individual genes (Figure 3B). Operon 1 contains *virB8-1 (DEJ70_04590), virB9-1* (*DEJ70_04585*), *virB10* (*DEJ70_04580*), *virB11* (*DEJ70_04575*) and *virD4* (*DEJ70_04570*). The vitamin B2 biosynthetic enzyme ribA encoded by *DEJ70_04595*, may be co-transcribed in this operon as observed in *w*Bm (Li & Carlow 2012). Operon 2 contains *virB3* (*DEJ70_01260*), *virB4* (*DEJ70_01265*), *virB6-1 (DEJ70_01270*), *virB6-2* (*DEJ70_01275*), *virB6-3* (*DEJ70_01280*) and *virB6-4 (DEJ70_01285*). Three duplicated genes: *virB4-2* (*DEJ70_01565*), *virB8-2* (*DEJ70_03190*) and *virB9-2* (*DEJ70_06825*) are found scattered elsewhere in the genome. Interestingly, *virB2* (*DEJ70_04445)*, which has been presumed absent from *Wolbachia* (Rancès et al. 2008), was also found in the *w*AlbB genome. Previous studies in other bacteria have shown that the T4SS is not constitutively expressed but tightly regulated by transcription factors (Félix et al. 2008), such as *wBmxR1* and *wBmxR2* in *w*Bm (Li & Carlow 2012). In *w*AlbB one corresponding homolog, *DEJ70_05760*, with higher sequence similarity to *wBmxR1* was found.

### Analysis of Ankyrin genes

Ankyrin repeat-containing (ANK) proteins are involved in protein–protein interactions and are rare in bacteria, but are found in *Wolbachia*, where they may be involved in host–*Wolbachia* interactions. The T4SS has been shown to be responsible for the secretion of the ankyrin repeat-containing protein AnkA in *Anaplasma phagocytophilum* (Lin et al. 2007), an intracellular bacterium closely related to *Wolbachia*. Based on genome-wide Pfam protein domain annotations (Table S1), 34 *w*AlbB proteins were found to contain at least one copy of an ankyrin repeat domain (Table 1), with a total of 81 copies of various types of ankyrin domains (Table S1). The same analysis performed for all complete *Wolbachia* genomes revealed a similar number of ANK proteins across insect *Wolbachia*, and fewer ANKs in filarial *Wolbachia* (Table 1).

### Identification of missing genes in *w*AlbB through KEGG pathway analysis

KAAS (KEGG Automatic Annotation Server) (Moriya et al. 2007) was used to obtain functional annotations of predicted protein sequences from the *w*AlbB genome. A total of 595 proteins were assigned a KO (KEGG Orthology) number. The KEGG pathway map and KO assignments for *w*AlbB were compared to those from the closely related *Wolbachia w*Pip and *w*Ri. Pairwise comparisons (*w*AlbB versus *w*Pip, *w*AlbB versus *w*Ri) of the KEGG annotated proteins revealed 5 proteins absent in *w*AlbB, namely DNA-3-methyladenine glycosylase (MPG, EC: 3.2.2.21), diacylglycerol kinase (DgkA, EC: 2.7.1.107), Cytochrome bd ubiquinol oxidase subunit I (CydA, EC: 1.10.3.14) and subunit II (CydB, EC: 1.10.3.14) and FtsI (EC: 3.4.6.4). The presence of these 5 proteins was determined in other *Wolbachia* genomes available in the KEGG database, namely *w*Mel, *w*Ri, *w*Ha, *w*No, *w*Pip, *w*Bm, *w*Oo and *w*Cle (Table 2). MPG was found in all other *Wolbachia* analyzed except in *w*AlbB, while the other 4 proteins were absent in *w*AlbB and in at least one additional species (Table 2).

**Table 2.**
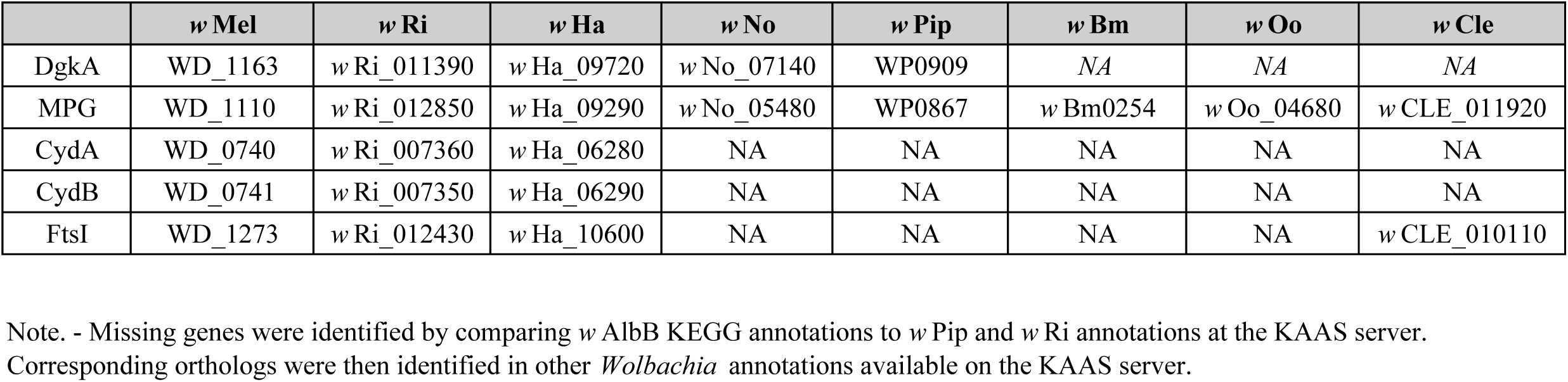
KAAS server based KEGG annotations identifies five genes missing in *w* AlbB.

## Discussion

We have assembled the complete genome of *w*AlbB from Aa23 cells. The key factors facilitating this were the relatively pure high molecular weight DNA extracted from host cell-free *Wolbachia*, and long read PacBio sequencing at high coverage.

The long reads enabled sequencing over complex repeat regions which have been difficult to resolve with short read sequencing. High depth of coverage (∼500X) enabled multiple rounds of polishing of the assembly to remove any SNPs or indel errors that are occasionally observed in technologies such as PacBio or Nanopore (Watson 2018). Absence of errors was also confirmed using Illumina data (∼600X coverage) generated from the same DNA sample.

*Wolbachia* genomes range in size from ∼0.9-1.8 Mb. Currently, 42 genomes have been deposited in the GenBank database, however, only 12 genomes show complete status. The complete circular *w*AlbB chromosome is 1,484,007 bp in size making it one of the largest sequenced *Wolbachia* genomes to date. The complete assembly is 321 kb larger than the draft sequence (Mavingui et al. 2012) and contains all the contigs from the published genome. Many of the gaps in the published draft were observed to be flanked by IS elements, suggesting that the repetitive nature of IS elements hinders assembly. Additionally, some contigs from the published draft mapped to multiple locations in the finished genome, indicating that they originate from repeated regions in the genome. Such repeated regions can be difficult to assemble using only short reads from Illumina or 454 platforms, but can be successfully assembled using PacBio long reads.

The genome of *w*AlbB contains 218 IS elements, belonging to 10 families, including one new family, scattered throughout the genome. The genomes of arthropod infecting *Wolbachia* have a large number of repetitive and mobile elements, particularly IS elements (Cerveau et al. 2011). A total of 11 distinct IS families was reported across *w*Bm, *w*Pel, *w*Ri, and *w*Mel genomes (Cerveau et al. 2011). The distribution and copy number of IS elements from different families varies between genomes (Cerveau et al. 2011). The *w*AlbB genome lacks members from IS6 and IS200/605 families, while IS982 (n=96) and IS481 (n=75) are present in higher abundance in comparison to other *Wolbachia* genomes (Cerveau et al. 2011). IS elements can cause disruption of genes, giving rise to pseudogenes. In *w*AlbB, many such examples were observed, e.g. pseudogenization of *rsmD* and *dgkA* genes.

Prophages represent another class of highly mobile elements that can have a significant impact on *Wolbachia* biology (Gavotte et al. 2007; Bordenstein & Bordenstein 2016; LePage et al. 2017; Beckmann et al. 2017). In a previous study of wild-caught, *Ae. albopictus* mosquitoes carrying either only *w*AlbA or both *w*AlbA and *w*AlbB, phage particles could be detected and quantified using qPCR (Chauvatcharin et al. 2006). In the current *w*AlbB genome, although 40 of the 63 genes from the WOVitA1 prophage (Bordenstein & Bordenstein 2016) could be found, many genes encoding essential components for a phage particle assembly were absent, while a few others were pseudogenized due to insertion of IS elements. In addition, the prophage related genes were found to be scattered over the *w*AlbB genome, unlike in an intact prophage.

Therefore the ancestral temperate prophage has undergone degradation in the *w*AlbB genome, a phenomenon also observed in other *Wolbachia* such as *w*Rec from *Drosophila recens* (Metcalf et al. 2014). It is therefore possible that the phage particles previously reported in *Ae*. *albopictus* mosquitoes (Chauvatcharin et al. 2006) were derived only from *w*AlbA, or the *w*AlbB phage was degraded during its *in vitro* culture in Aa23 cells. Interestingly, orthologs of the 2 WO prophage proteins cifA and cifB involved in cytoplasmic incompatibility (LePage et al. 2017; Beckmann et al. 2017) are encoded in *w*AlbB by two pairs of genes, but are not in close proximity to the prophage-derived genes or gene clusters. These gene-pairs might be remnants from an earlier integrated prophage which has since undergone degradation. Similarly, in another *Wolbachia, w*Rec, approximately 75% of its prophage gene content has been lost (Metcalf et al. 2014), but still retains an intact *cifA, cifB* gene pair (Lindsey et al. 2018).

Orthology analysis of all annotated proteins from *w*AlbB and 12 other complete *Wolbachia* genomes identified the core *Wolbachia* proteome comprising 536 orthogroups. The majority of these (n=519) contain single-copy, 1:1 orthologs which are ideally suited for phylogenomic analysis. Further analysis of this core proteome could shed light on the unique intracellular lifestyle of *Wolbachia* and provide insight into essential *Wolbachia* genes that may be targeted in an anti-symbiotic approach to new drug discovery in filarial parasites. On the other hand, studies on orthogroups unique to a particular genome (e.g. 13 proteins present only in *w*AlbB) may lead to the discovery of proteins which are involved in the adaptation of *Wolbachia* to a particular host/cell niche.

The T4SS of *w*AlbB is encoded by two operons and a few genes scattered throughout the genome. Their organization and sequence are conserved across various *Wolbachia*, likely due to their important roles in its biology, such as secretion of effectors that influence host:bacteria interactions. Several candidate effectors of the T4SS in *w*Mel were identified recently (Rice et al. 2017), including one which interacts with the host cytoskeleton (Sheehan et al. 2016). Ankyrins are established T4SS effectors of intracellular bacteria. The genome of *w*AlbB encodes 34 ANK proteins. *ANK* genes in *w*Pip are linked to polymorphisms in cytoplasmic incompatibility phenotypes in *Cules pipiens* mosquitoes (Sinkins et al. 2005). In the closely related *Anaplasma* and *Ehrlichia*, an ankyrin repeat-containing (ANK) protein has been shown to regulate transcription, suppress host innate immunity, inhibit host cell apoptosis and reduce reactive oxygen species (Rikihisa & Lin 2010; Liu et al. 2012). However, most *ANK* genes contain many copies of short open reading frames of unknown function (Wu et al. 2004). The T4SS could also be involved in lateral gene transfer events between *Wolbachia* and its host (Dunning Hotopp et al. 2007). For example, VirB6, an essential trans-membrane channel component of the T4SS in many bacteria, has been shown to direct DNA export through the T4SS in *Agrobacterium tumefaciens* (Jakubowski et al. 2004). Interestingly, all *Wolbachia* genomes, including *w*AlbB, encode 4 VirB6 paralogs.

Comparing the KEGG pathway maps and KO assignments in *w*AlbB with those in closely related *w*Pip and *w*Ri identified 5 proteins that were absent in *w*AlbB, namely DgkA, MPG, CydA, CydB and FtsI. In the *w*AlbB genome, *dgkA* is pseudogenized due to insertion of an IS982 family transposase, while the gene is intact in *w*Mel, *w*Ri, *w*Ha, *w*No and *w*Pip, but is absent in *w*Bm, *w*Oo and *w*Cle. DgkA phosphorylates diacylglycerol to generate phosphatidic acid in glycerolipid and glycerophospholipid metabolism pathways and plays an important role in microbial stress responses (Yamashita et al. 1993). MPG is involved in the DNA base excision repair pathway by recognizing a variety of base lesions, mainly caused by alkylating agents, resulting in release of the damaged base in free form from alkylated DNA and initiation of repair (Costa de Oliveira et al. 1987). *MPG* is present in all the *Wolbachia* examined here, except *w*AlbB. The genes *cydA* and *cydB* are present in *w*Mel, *w*Ri, *w*Ha, and *w*No, yet absent in *w*Bm, *w*Oo, *w*Cle and *w*AlbB. The proteins CydA and CydB are members of a family of integral membrane proteins involved in catalyzing terminal electron transfer in eubacterial and archaeal respiration. Their high oxygen affinity enable them to scavenge and reduce oxygen to water, preventing damage to oxygen-sensitive enzymes, and permitting growth in microaerobic and anaerobic environments and survival under a number of stress conditions (Borisov et al. 2011). FtsI is a class B3 penicillin-binding protein (PBP B3) that functions as a transpeptidase in peptidoglycan metabolism, essential for bacterial cell wall synthesis and cell division (Cayô et al. 2011). Although *w*AlbB does not have FtsI, it does contain a class PBP B2 transpeptidase, MrdA, while other *Wolbachia, w*Mel, *w*Ri, *w*Ha, and *w*Cle, have both FtsI and MrdA, indicating potential redundancy.

The availability of a complete circular genome from *w*AlbB will provide further insight into phylogenetic relationships between the different *Wolbachia* supergroups, and enable further biochemical, molecular and genetic analyses on *w*AlbB and related *Wolbachia*. The annotation and analysis of mobile elements highlight their considerable effect on genome evolution and gene content in intracellular symbionts, suggesting that such elements could be re-purposed as tools for genetic manipulation of *Wolbachia.* The genome also provides an important baseline for further studies of *Wolbachia* interactions with its host that may advance practical applications such as the use of *Wolbachia* for pest and disease control.

## Acknowledgments

The authors thank the following for helpful discussions and comments on the manuscript: Brian Anton, Rich Roberts, Peter Weigele, Rick Morgan, Tom Evans, Bill Jack, Jeremy Foster, Barton Slatko, Emilie Lefoulon, Youseuf Suliman and Catherine Poole; and the DNA sequencing core at New England Biolabs for Illumina sequencing. The authors are also grateful for the continued encouragement from Don Comb. This work was supported by New England Biolabs.

## Figure Legends

**Fig. S1.**
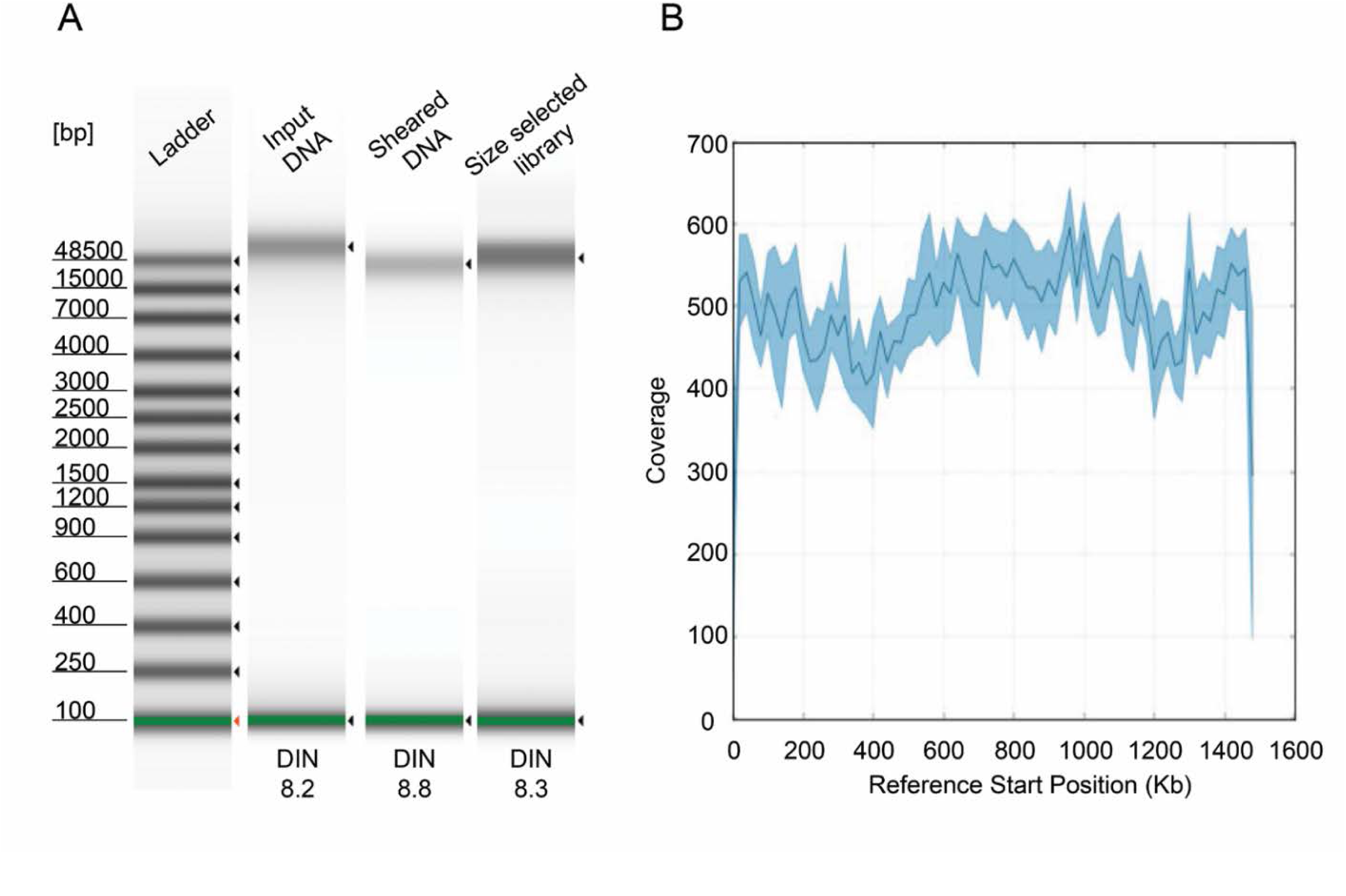
PacBio library containing large inserts yields *w*AlbB draft contig with ∼500X coverage. (A) Gel image of genomic *w*AlbB DNA and PacBio library analyzed using the Agilent 4200 TapeStation system. (B) Coverage report of *w*AlbB draft contig produced by first round of PacBio HGAP3 assembly.

**Fig. S2.**
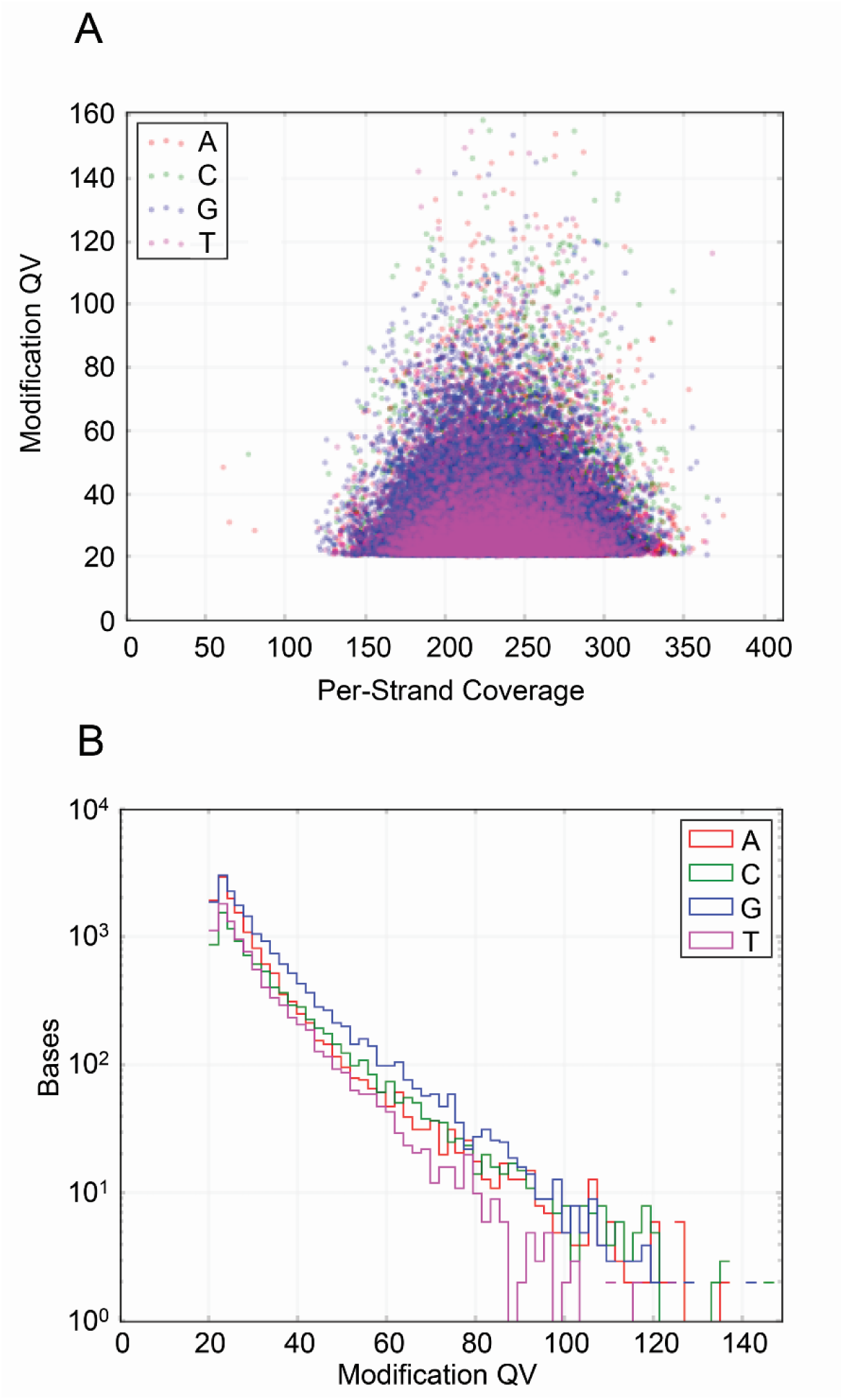
No detectable DNA methylation in *w*AlbB genome. Using the PacBio RS_Modificaton_and_Motif_Analysis.1 pipeline, no robust signal for DNA modifications such as m6A and m4C in *w*AlbB genome were detected. (A) The “Modification QV” quality scores for each of the 4 bases A, C, G and T were almost normally distributed over the entire coverage range, unlike genuine modifications where the scores increase lineary with covrage (See https://github.com/PacificBiosciences/Bioinformatics-Training/wiki/Methylome-Analysis-Technical-Note)

**Fig. S3.**
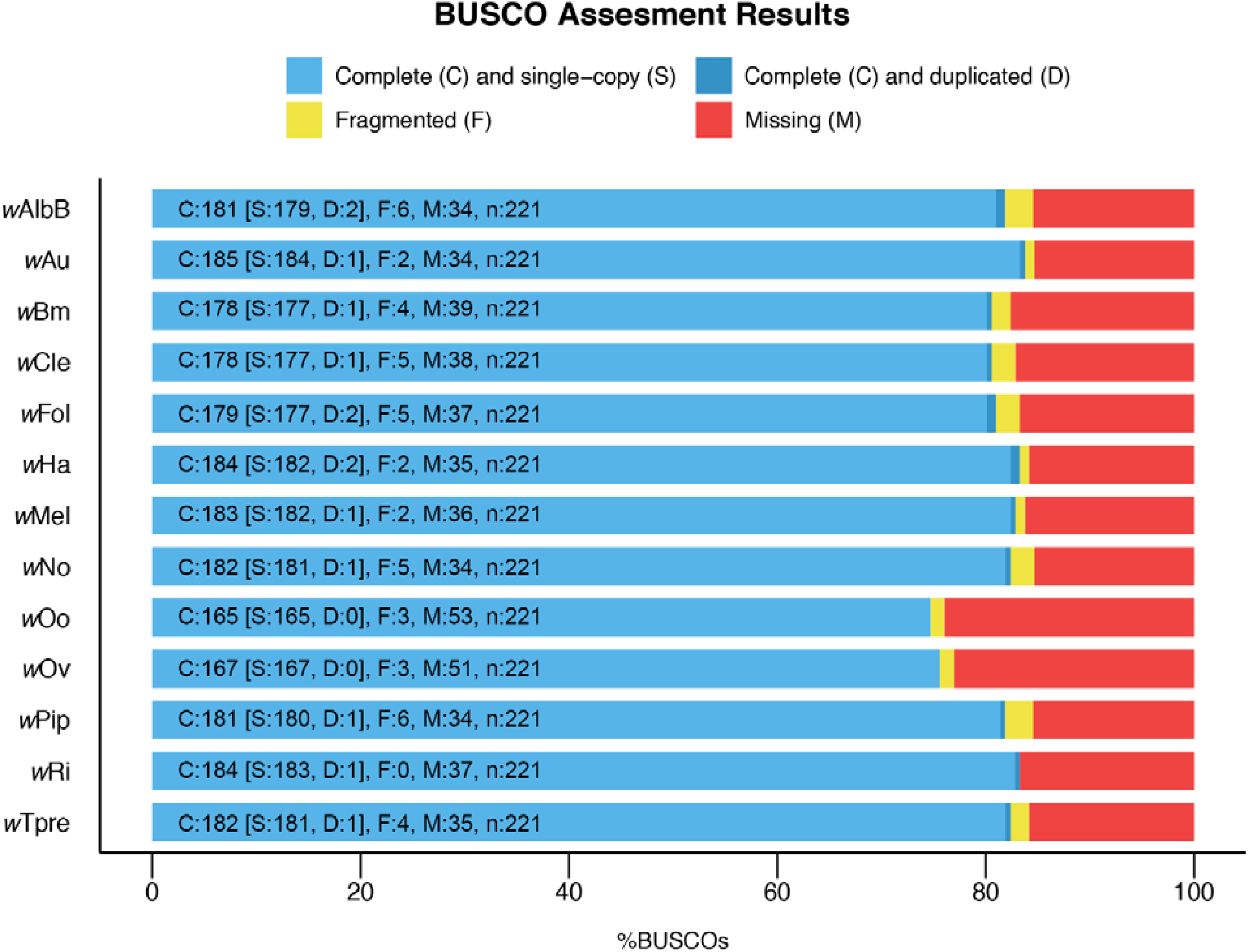
Similar BUSCO scores across all the complete *Wolbachia* genomes. The BUSCO pipeline was used to measure the proportion of highly conserved, single-copy orthologs (BUSCO groups). The set of reference BUSCO groups was set to the lineage “Proteobacteria”, which contains 221 BUSCOs derived from 1,520 proteobacterial species.

